# Dissecting individual pathogen-commensal interactions within a complex gut microbiota community

**DOI:** 10.1101/2020.04.06.027128

**Authors:** Jack Hassall, Meera Unnikrishnan

## Abstract

Interactions of commensal bacteria within the gut microbiota and with invading pathogens are critical in determining the outcome of an infection. While murine studies have been valuable, we lack *in vitro* tools to monitor community responses to pathogens at a single-species level. We have developed a multi-species community of nine representative gut species cultured together as a mixed biofilm and tracked numbers of individual species over time using a qPCR-based approach. Introduction of the major nosocomial gut pathogen, *Clostridiodes difficile*, to this community resulted in increased adhesion of commensals and inhibition of *C. difficile* multiplication. Interestingly, we observed an increase in individual *Bacteroides* species accompanying the inhibition of *C. difficile*. Furthermore, *Bacteroides dorei* reduced *C. difficile* growth within biofilms, suggesting a role for *Bacteroides spp* in prevention of *C. difficile* colonisation. We report here an *in vitro* tool with excellent applications for investigating bacterial interactions within a complex community.

## Introduction

The gut microbiota, which is the largest microbial community found in the human body, plays a key role in an array of essential physiological processes, including immune function, metabolism and nutrient absorption. Imbalances and shifts in the gut microbiota composition have been associated with multiple conditions including chronic gastrointestinal diseases like inflammatory bowel disease (IBD)^1,2^ and systemic metabolic diseases like diabetes and obesity^3,4,5,6^. Most studies linking disease states to the microbiome are based on 16S rRNA or whole microbial genome sequencing, although recent studies have begun to demonstrate several interesting mechanisms underlying microbiota functions^7^. An important function of the gut microbiota is to also form a protective barrier against colonisation by gastrointestinal pathogens. a property described as colonisation resistance^8,9^.

Colonisation resistance occurs through an array of direct or indirect bacterial and host interactions including competition for nutrients, host metabolites and physical space^10^. An example of nutrient competition is the commensal *Bacteroides thetaiotaomicron* which consumes carbohydrates essential to murine pathogen *Citrobacter rodentium* causing it to be excluded^11^. Secreted compounds released by the microbiota such as the antimicrobial peptides, bacteriocins and short chain fatty acids (SCFA) can directly affect an invading pathogen. *Bacteroides* were shown to inhibit *Salmonella Typhimurium* through the production of the SCFA, propionate^12^. SCFAs also impact epithelial barrier function by affecting production of host molecules including antimicrobial peptides and epithelial mucins^13,14^. Disturbed microbiota and the loss of colonisation resistance are associated with several pathogen infections including *Clostridioides difficile^8^, Enterohemorrhagic E. coli*^15^, and *Campylobacter jejuni*^16^.

In the case of *C. difficile*, a leading cause of healthcare-associated diarrhoea worldwide, colonisation occurs only when the microbiota is altered, usually due to treatment with antibiotics such as fluoroquinolones^17^. The increased susceptibility of *C. difficile* infection (CDI) after antibiotic-induced dysbiosis of the gut microbiota is well documented^18,19^. While most studies demonstrating the link between CDI and antibiotic therapy are based on changes in microbial populations by microbiota sequencing, recent studies have reported mechanisms by which the microbiota can prevent *C. difficile* infections^20^. The microbiota in a healthy state consumes or converts primary bile acid into secondary bile acids, reducing the ability of *C. difficile* to germinate^21^. Secondary bile acids such as deoxycholic acid (DCA) and lithocholic acid (LCA), are toxic to vegetative *C. difficile*^22,23^. Additionally, gut bacteria like *Clostridium scindens* which encode secondary bile acid synthesis enzymes have been associated to resistance to *C. difficile* infection^24^. The microbiota not only competes for resources but actively inhibits *C. difficile* through production of bacteriocins such as the thuricin CD, produced by *Bacteroides thuringiensis*^25^.

While the microbiota is clearly important in preventing infections, current knowledge is mainly based on microbiota profiles from faeces. The gut microbiota is composed of bacteria within the lumen, which are usually detected in faeces and bacteria associated with the gut mucosa. Few studies have profiled the adherent microbiota population of healthy human guts as they require invasive biopsies^26^. While not much is known about the composition and dynamics of this population it can be viewed as a mixed biofilm community that is closely associated with mucus layers in the gut, which provide a spatial and metabolic niche for the bacteria^27,28^. The understanding of how individual bacteria within such complex communities interact remains poorly understood. *In vitro* systems that mimic gut microbial communities and are easily trackable are necessary to study inter-bacterial interactions.

Identifying and quantifying species in a mixed community is challenging, simple microscopy cannot be used as cells are often morphologically too similar. Selective media have proven to be successful with small communities, however the difficulty of finding species-specific media increases as the community gets larger^29^. Quantitative PCR (qPCR)-based approaches have been shown to be successful at predicting cell concentration and biomass formation of individual species in mixed populations^30,31^. qPCR has been routinely used for accurate quantification of oral pathogenic species like *Streptococcus mutans* and *Streptococcus sobrinus* from dental biofilms in order to help decide treatment effectiveness^32,33^. qPCR has also been widely used to quantify intestinal bacterial species in faeces^34,35^.

In this study, we create a representative adherent multi-species gut community, in which we can track behaviours of individual species over time using a qPCR-based method. We have employed this system to study how individual commensal species behave in the presence of the human pathogen *C. difficile*. We report an increase in *Bacteriodes spp* within this complex biofilm community and a direct impact of a *Bacteroides* species on the growth of *C. difficile*.

## Methods

### Bacterial culture

All the bacterial strains listed in Table 1 were cultured at 37°C in anaerobic conditions using an anaerobic cabinet (Don Whitley DG250), and unless stated otherwise cultures were grown in Schaelder Anaerobic Broth (SAB, Oxoid) supplemented with 0.005 mg/ml Vitamin K (VWR) and 2 mg/ml L-cysteine (Sigma-Aldrich) (Referred to now as SAB+).

**Table 1:** List of representative species used to construct a gut microbial community.

| Species | Phylum | Source |
| --- | --- | --- |
| *Bacteroides dorei* (CL02T00C15) | Bacteroidetes | BEI resources |
| *Bacteroides ovatus* (3_8_47FAA) | Bacteroidetes | BEI resources |
| *Bacteroides thetaiotaomicron* (VPI-5482) | Bacteroidetes | Dr Anne Marie Krachler |
| *Bifidobacterium adolescentis* (L2-32) | Actinobacteria | BEI resources |
| *Blautia hansenii* (20583) | Firmicutes | DSMZ |
| *Clostridioides difficile* (R20291) | Firmicutes | Dr Trevor Lawley |
| *Escherichia coli* (83972) | Proteobacteria | BEI resources |
| *Eubacterium hallii* (3353) | Firmicutes | DSMZ |
| *Faecalibacterium prausnitzii* (17677 A2-165) | Firmicutes | DSMZ |
| *Ruminococcus gnavus* (CC55_001C) | Firmicutes | BEI resources |

DSMZ: Deutsche Sammlung von Mikroorganismen und Zellkulturen and BEI: Biodefense and Emerging Infections Research Resources Repository.

### Microbiota biofilm assay

For mixed biofilms with multiple microbiota species, individual bacterial cultures were grown in SAB+ for 16-18 h (overnight at 37°C in an anaerobic cabinet). These cultures were then diluted with fresh SAB+ to achieve a final concentration of 0.1 OD600 per species within the mixed culture. To ensure consistency in experiments with differing numbers of species (with and without *C. difficile)*, we supplemented the absence of a species with an equivalent amount of medium, i.e. in each condition the concentration of the conserved species remained the same. The diluted cultures were added together and inverted several times to ensure a homogenous mix. 1 ml of culture mixture was added per well of a 24-well polystyrene tissue culture treated dish and biofilms were allowed to form at 37°C for the required time (6 h-72 h) in an anaerobic cabinet. At each timepoint the wells were gently washed twice with 1 ml PBS and resuspended in PBS. The resuspended biofilms were then treated with PMA, after which a DNA extraction was conducted.

### Propidium monoazide treatment

Prior to DNA extraction, propidium monoazide (PMA, Biotium) was added to the resuspended samples at a final concentration of 160 μM. The samples were then incubated in the dark for 10 min (37°C, in anaerobic cabinet). To photoactivate the PMA, samples were activated in the PhAST Blue (Geniul) light system for 15 min, following which genomic DNA was extracted.

### Genomic DNA extraction

DNA extraction was carried out using a phenol chloroform-based method. Cultures were centrifuged at 14,000 rpm for 5 min, after which the supernatant was discarded. The cell pellets were re-suspended in 500 μl of 5 mg/ml lysozyme (VWR) and incubated for 20 min at 37°C. Following this, 20 μl RNase solution (20 mg/ml) (Fisher Scientific), 20 μl proteinase K (20 mg/ml) (New England Biolabs) and 25 μl 10sSodium dodecyl sulphate (SDS) (Fisher Scientific) were added. The samples were then incubated at 37°C for a further 10 min, following this 100 μl of NaCl (Fisher Scientific) and 80 μl Cetyl Trimethyl Ammonium Bromide (CTAB) (Sigma Aldrich) were added before a final incubation at 60°C for 45 min.

The lysed samples were treated with 750 μl of phenol chloroform IAA (PCI) (Sigma Aldrich), vortexed (2-3 sec) and centrifuged for 10 min at 14,000 rpm. The upper phase was transferred to a fresh Eppendorf tube, and the PCI centrifuge steps were repeated until a clear boundary could be seen between the two phases. Next, 75 μl of 3M sodium acetate (pH 5.2) (Sigma Aldrich) and 750 μl of cold (−20°C) 96% ethanol were added. This solution was inverted until the DNA precipitated out. The precipitated DNA solutions were then centrifuged for 5 min at 14,000 rpm. The pellets were washed with 200 μl 70% ethanol and centrifuged for a final time (2 min at 14,000 rpm). The DNA pellet was dried at room temperature and re-suspended in 75 μl TE buffer.

### Real time quantitative PCR (qPCR)

Primers were designed to target either the topoisomerase I (*topI*) or DNA gyrase subunit A (*gyrA*) region of each strain (Supp Table 1 for sequences). The primers were designed using Primer-BLAST, this enables a high level of species specificity to be engineered *in situ*^36,37^. Primers were designed to create an amplicon of 100-150 bases, with an annealing temperature of 59-60°C, all primers were screened against the ‘nt’ database for specificity. Suitable primers were then checked using ‘Thermo Fisher’s Multiple Primer Analyzer’, a web-based script which looks for any possible primer-dimer structures. Any primers found to form dimers were ruled out. The qPCR was conducted with the Agilent Mx3005P qPCR System and the Luna^®^ Universal qPCR Master Mix (New England Biolabs). The temperature profile used throughout was the recommended standard for the master mix.

To convert from a qPCR output cycle threshold (Ct) to a predicted bacterial number we utilized previously described methods^32,37^. This required creating a standard curve for each species, the gradient of which is related to the primer amplification efficiency. The standard curve is a serially diluted sample where the Ct value and the DNA mass for each dilution is known, and the curve maps the relationship between the two. DNA concentration was measured using the Qubit fluorometer 2.0 (Thermo Fisher) and the dsDNA Qubit kit (Thermo Fisher), depending on the mass this would vary between high sensitivity and broad spectrum.

Once the standard curve was established, we used it to convert Ct values into the starting mass of DNA (M_DNA_) (Equation 1). Using the assumption that one genome weight worth of DNA is equal to one bacterium, we were able to predict the total number of bacteria by dividing the starting amount of DNA by the calculated genome mass (MGenome) (Equation 2). The mass of each bacterial genome was calculated by multiplying the length of the genome (GLength) by the average weight of one base pair (bp). Commonly the average weight of a nucleotide is expressed as the weight of one mole (WBase) not a singular base, this is equal to 650 da. To convert from one mole to 1 bp we divided through by Avogadro’s number (N_A_).

Equation 1: Using the gradient (m) and intercept (c) from the standard curve, we can use the cycle threshold (CT) value of the current qPCR run to predict the starting DNA amount (M_DNA_).

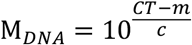

Equation 2: Assuming one bacterial weights worth of DNA is equal to one bacterium we can divide the predicted starting genome mass (M_DNA_) by the mass of one bacteria genome (M_Genome_) to get an estimate on the bacterial number. The mass of one genome is calculated from the genome length (G_Length_) and the average weight of one mole of nucleotide bases (W_Base_). Avagadro’s number (N_A_) is used to convert average weight of one mole to the average weight of a base pair.

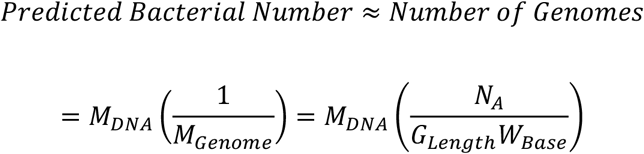

### *C. difficile* and *B. dorei* biofilm studies

*B. dorei* and *C. difficile* were cultured overnight in brain heart infusion (BHI) media (Sigma Aldrich) supplemented with 0.5 mg/ml yeast extract (Fisher Scientific) and 0.001 mg/ml L-cysteine (BHIS)^38^. After which, cultures were diluted down to 0.1 OD and added to a 24-well polystyrene tissue culture treated plate. Each well was made up to a total of 1 ml with monocultures having a mix of 0.5 ml culture and 0.5 ml BHIS media, and cocultures containing 0.5 ml of each species. The resulting end concentration was 0.05 OD_600_. At set timepoints the biofilms were washed twice with 1 ml PBS, then manually resuspended in 1 ml PBS. Dilutions were plated on BHIS with *C. difficile* supplement (Oxoid). *C. difficile* and *B. dorei* have two distinct colony morphologies^38^, which allowed us to differentiate between *Bacteroides* and *C. difficile* and quantitate each species

### Statistical analysis

All experiments were performed in triplicates and repeated at least three times independently. An unpaired Student’s t-test was used to determine if differences between two groups were significant, and a two-way ANOVA was used to compare multiple groups.

## Results

### Developing a mixed biofilm community and optimisation of a PMA-qPCR method for tracking individual species

In order to develop a complex mixed biofilm community comprising of representative gut species, we selected a total of nine gut commensal species were selected (Table 1), based on previous available literature describing species present in healthy human gut microbiota^3,39–41^. The species were selected based on high relative abundance, presence across multiple regions (Europe, America and Asia), and the availability of a sequenced genome. *Firmicutes* and *Bacteroidetes* were overrepresented as these genera are the dominant phylum in the gut^42^. These nine species were cultured together as a mixed biofilm as described in Methods.

In order to track individual species, primers were designed to target either the topoisomerase I (*topI*) or DNA gyrase subunit A (*gyrA*) region of each strain. These genes were chosen because unlike the conventional 16S gene, they have a single copy number within the genome which improves the accuracy of assumptions made in converting DNA mass to bacteria number. For qPCR-based quantification to be successful primers have to be highly specific, with little to preferably no off-target amplification. To test specificity, we tested genomic DNA obtained from each species against primers specific to all the species. We found high levels of specificity with no cross-reacting bands outside of the correct lane (Supp Figure 1).

For accurate quantification of bacteria, it is essential to quantify only active/live cells. Standard qPCR quantifies all cells, ‘dead’ and ‘active’ alike, which gives an inaccurate abundance of individual species within a population. To improve the accuracy of our quantification we used propidium monoazide (PMA) to prevent the counting of ‘dead’ bacteria, i.e. those with compromised membranes. PMA is a photo-activated DNA binding dye that can only target cells with ruptured membranes or ‘dead’ cells^43–46^. When PMA is photo-activated, it covalently binds to dsDNA, crosslinking the two strands, and this binding prevents DNA amplification during PCR^33^. Hence, biofilm samples are first treated with an optimised concentration of PMA, followed by genomic DNA extraction and then analysis by qPCR (Figure 1).

**Figure 1:**
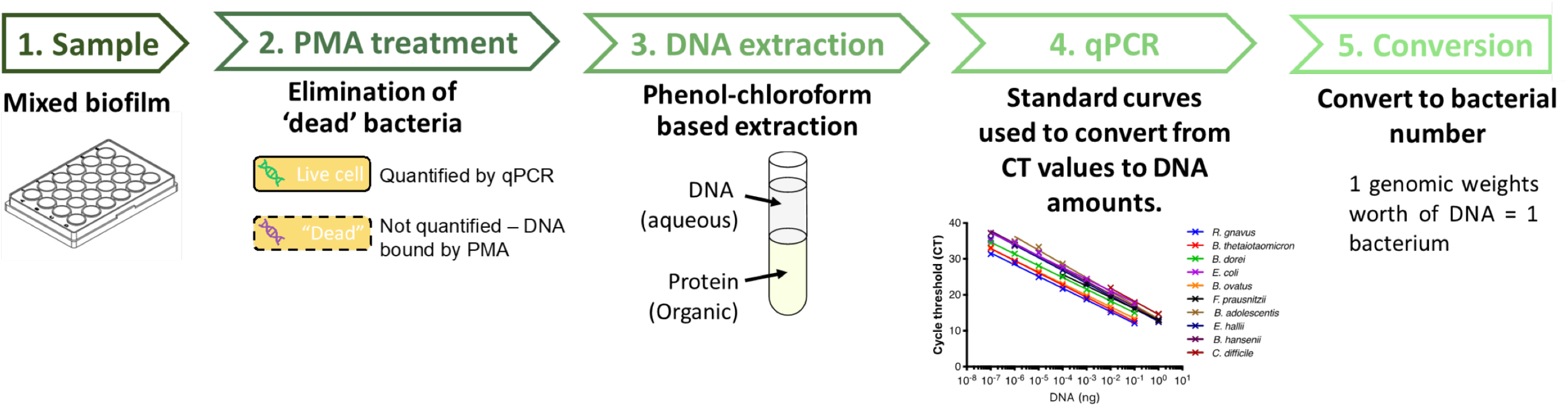
Schematic diagram of the pipeline used to quantify individual species in a mixed biofilm. Samples are PMA (propidium monoazide) treated to ensure quantification of genomic DNA only from live cells. After treatment, cells are lysed, DNA is extracted, followed by qPCR quantification of DNA and subsequent conversion to bacterial numbers.

To convert the qPCR cycle threshold (Ct) values to a DNA mass, a standard curve for each species was generated (Supp Figure 2). The standard curve consists of the Ct value plotted against the known DNA mass (measured by Qubit) of serially diluted DNA samples. A semilog line was fitted to each curve, with a regression above 0.990 for all (Supp Table 2). The primer efficiency for each species was calculated from the regression. An efficiency between 90 and 110% is generally accepted as good enough for accurate qPCR^47^, and our chosen species all fall in this range except for *B. adolescentis* and *C. difficile*. Given that both are above 80% efficient and linear (semilog) over the range of DNA values we tested, we believe they are acceptable for the demands of our experiments. Once we converted Ct values into a DNA mass using the standard curve, we then converted it finally into a bacterial number. To achieve this, we used the assumption that one genomic weight worth of DNA is equal to one bacterium. We established genome weight using genome sequence length and the average weight of a single base pair (Supp Table 3).

To confirm the accuracy of this qPCR-based quantitation assay, we grew single species biofilms and tested the prediction bacterial number from qPCR versus the actual CFU values obtained from plating (Supp Figure 3). We chose a species representative of each phylum (Firmicutes, Bacteroidetes, Proteobacteria and Actinobacteria). We found that the predicted values are similar to that of the CFU assay, a two-way ANOVA predicts there is no significant difference between the two techniques.

### Tracking individual species within a microbiota community

We first tracked numbers of individual species within a mixed biofilm containing nine species. All nine species were detected after different times of incubation up to 72 h (Figure 2B). *Bifidobacterium adolescentis* and *B. ovatus* both showed a positive trend at early timepoints (before 72 h), while *B. thetaiotaumicron* remained unchanged and all other species showed a reduction in numbers. At 72 h, all the species showed a decay in numbers, implying a build-up of toxic secreted bi-products in the medium, or nutritional depletion. Lower bacterial numbers for some species (*Blautia hansenii*, *Bacteroides thetaiotaumicron, Bacteroides ovatus* and *Bacteroides dorei*) in the inoculum (Supp Figure 4) (in spite of normalising numbers by optical density), appeared to impact their numbers within the mixed biofilm. Our results show that species within the biofilm, even within the same genus, showed distinct behaviours, with *Bacteroides* having positive, neutral and negative trends over the first 48h.

**Figure 2:**
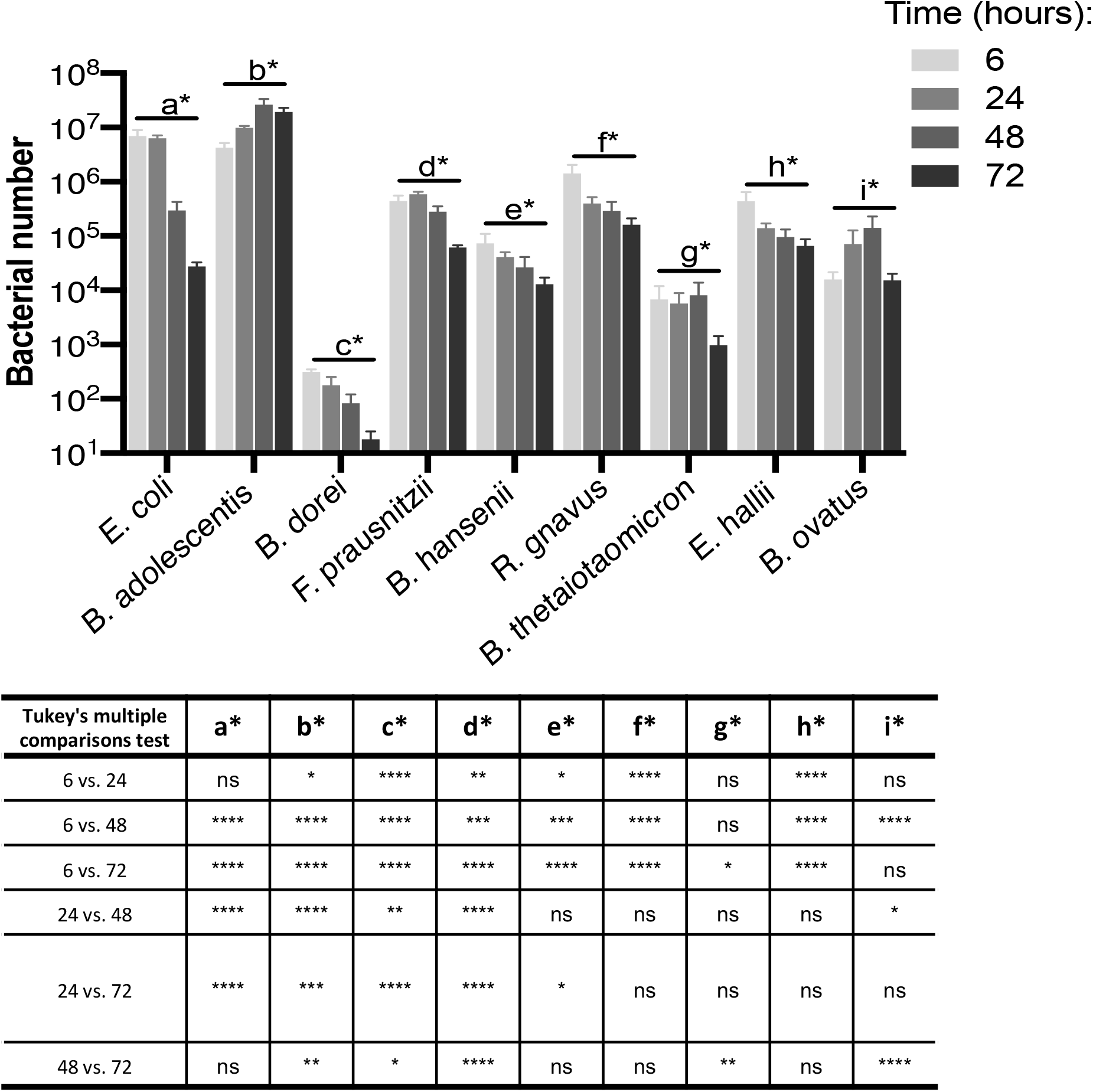
A nine-species gut microbial community tracked over time. Species-specific changes in bacterial numbers within a mixed biofilm were tracked using PMA-qPCR. Ct values were converted using our pipeline to represent total bacterial numbers. Bacterial numbers for individual species within the mixed biofilm at 6, 24, 48 and 72 h. The table below summarises the significant differences across time for the different species as calculated by ANOVA, with post hoc Tukeys’ multuiple comparison test. Data shown are the mean of three independent biological experiments done in in triplicate. Error bars indicate the standard deviation.

### Microbiota species interfere in *C. difficile* adherence and growth

In order to study the effect of a gut pathogen on the dynamics of this gut microbiota community, we chose to study the effects of the nosocomial pathogen *C. difficile*. First, to study the effects of the microbiota on *C. difficile* adhesion, we tracked the formation of adherent biofilms over 6 h in the presence and absence of *C. difficile*. We measured the percentage of the initial inoculum that is able to adhere to a 24-well polystyrene plate. A significant reduction in initial adhesion was observed in *C. difficile* when cultured with the microbiota compared to a *C. difficile* monoculture control (Figure 3A**Error! Reference source not found.**). Although significant, this reduction is small, with ~8% of the initial inoculum adhering when cultured alone and ~4% with the microbiota. With respect to the microbiota, we saw that when cultured alongside *C. difficile* there is a significant increase in the number of bacteria that adhered. Statistically significant differences were seen for *Escherichia coli, B. adolescentis* and *Ruminococcus gnavus*, but the trend can be seen generally across all species (Figure 3B). *E. coli* appeared to be dominating at this early stage with far more of its original inoculum adhering than any other species. *E. coli* was the only facultative species present so any lingering oxygen in the reduced media may provide it with an initial head start.

**Figure 3:**
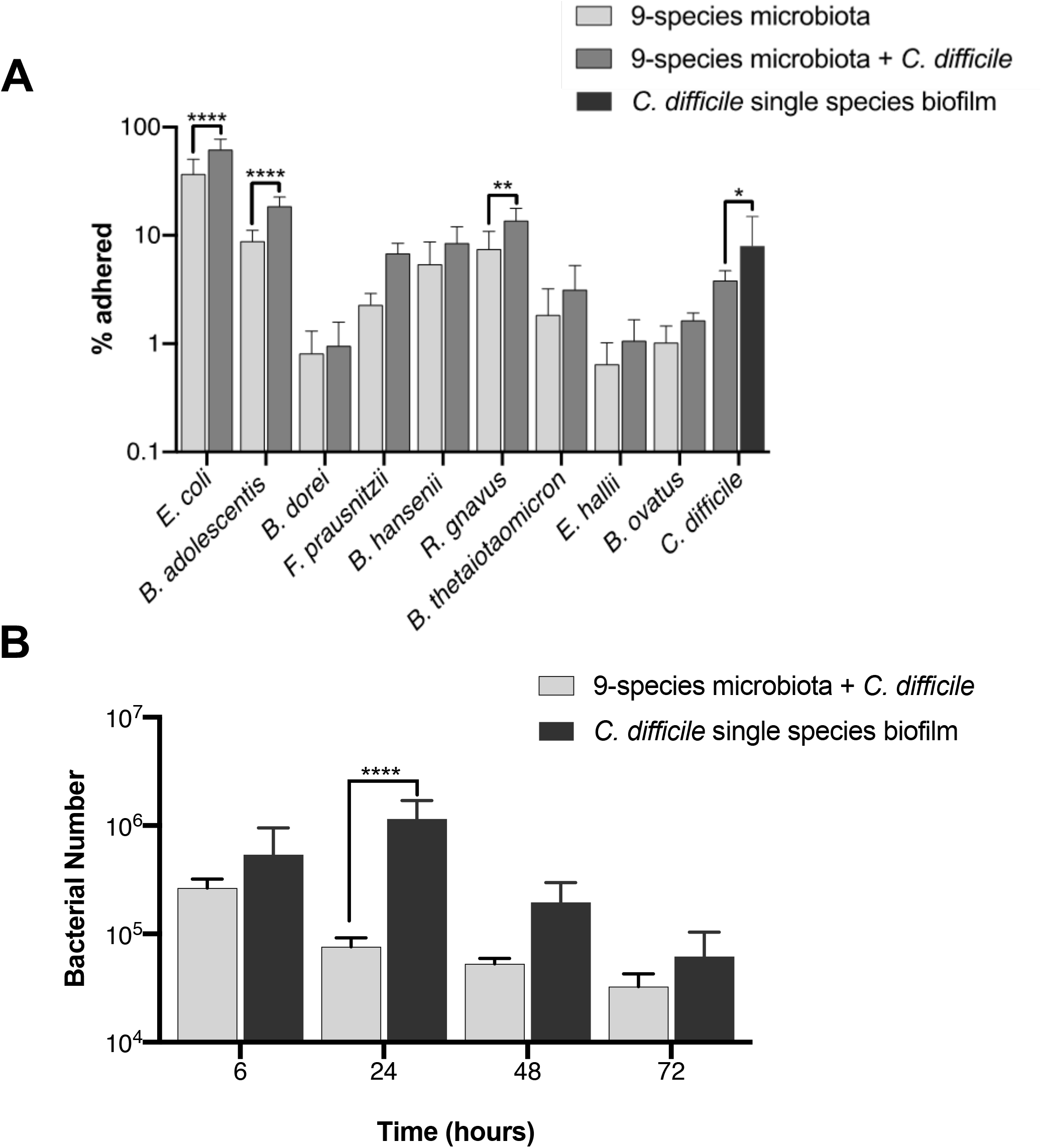
Interactions of *C. difficile* with a commensal microbiota community. **A)** The presence of *C. difficile* impacts the adhesion of several species in the adherent microbiota community. The percentage of the inoculum which adhered after 6 h, in a nine species microbiota community (9-species microbiota), in the nine species community with *C. difficile* (9-species microbiota + *C. difficile*), and a single species *C. difficile* biofilm control (*C. difficile* only biofilm). **B)** The microbiota has an inhibitory effect on *C. difficile. C. difficile* bacterial numbers when cultured in monoculture biofilm or with a nine species representative microbiota were tracked over 72 h using PMA-qPCR. All data shown are of three independent experiments in triplicate. A two-way ANOVA indicates a significant difference between the two conditions (*P*-value < 0.0001), with the post-hoc Sidak’s test used to determine specific difference, *P*-values - **** < 0.0001, ** < 0.01, * < 0.05.

We next investigated the impact on *C. difficile* during biofilm formation (Figure 3B). Here the microbiota impacts *C. difficile* growth, significantly reducing it at all time points when compared to a *C. difficile* monoculture control. This impact was the highest at 24 h, with the microbiota causing a 10-fold drop in *C. difficile* numbers compared to the monoculture control. Given the higher inhibition seen post-adhesion, it is likely that the decrease in *C. difficile* numbers is likely attributed to the microbiota negatively impacting *C. difficile* growth, rather than a reduction in the ability of *C. difficile* to adhere.

In the presence of *C. difficile*, we tracked each of the nine species that make up the representative microbiota (Figure 4 and Supp Figure 5). When comparing this to a microbiota only control, we see that the presence of *C. difficile* has a neutral to positive effect, with five of the nine species having a small but significant difference for *B. dorei, B. ovatus, E. coli, Faecalibacterium prausnitzii* and *R. gnavus* at 6 h and/or at 24 h and 48 h. Although *C. difficile* increases initial binding in the microbiota, the effect is not seen long term. In both conditions, all species no matter the prior trajectory, show a decrease in numbers at 72 h. A lack of sufficient nutrients in the medium or the accumulation of toxic levels of secreted bi-products at this late time point could negatively impact bacterial growth.

**Figure 4:**
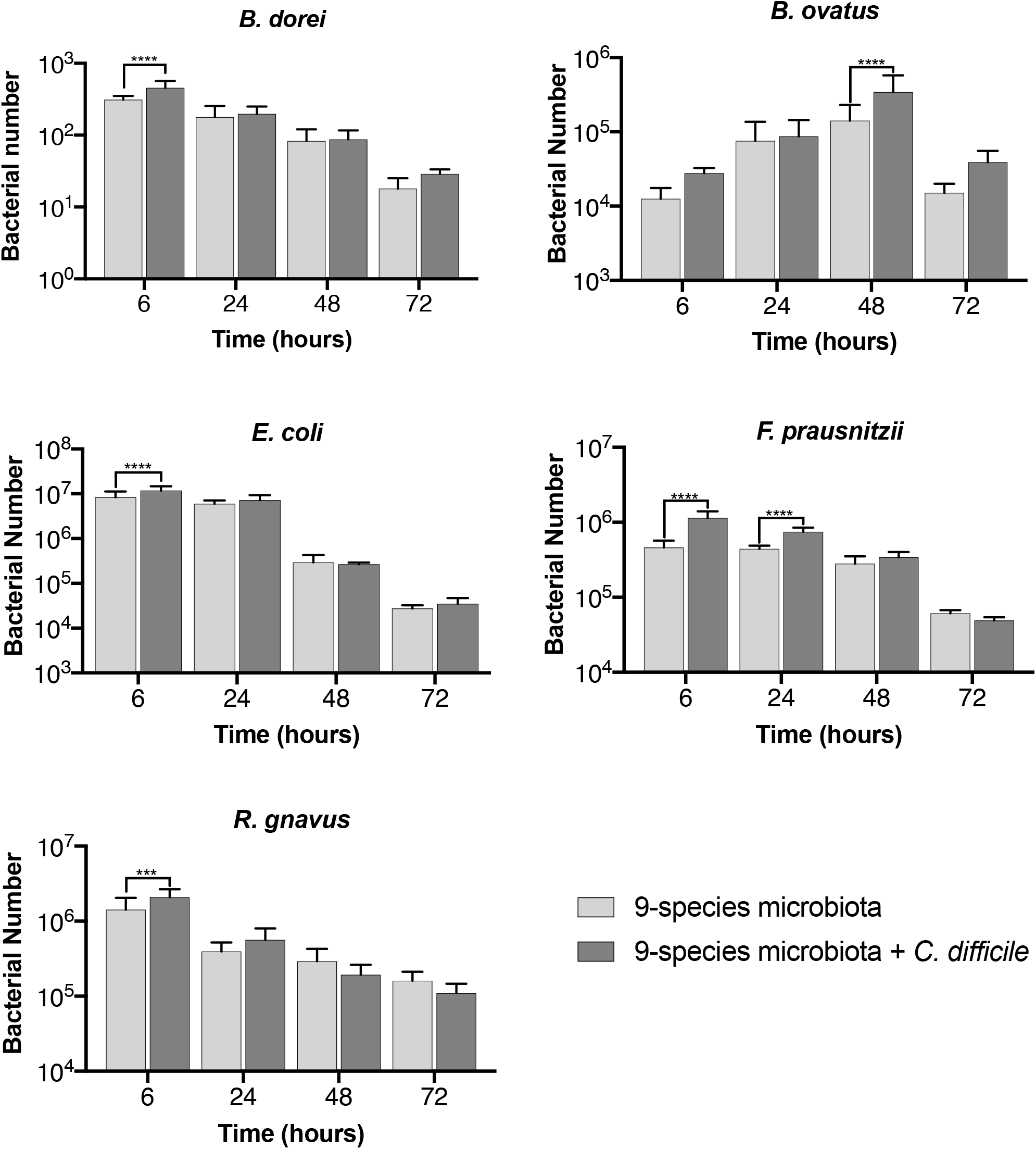
Tracking the effects of *C. difficile* on individual species within the gut microbiota community. Bacterial numbers of individual species within a representative microbiota and *C. difficile* biofilm over 72 h using PMA-qPCR are shown compared to a control microbiota biofilm without *C. difficile*. Data shown are of three independent experiments in triplicate. A two-way ANOVA was used to determine significant difference between the two conditions, a post-hoc Sidak’s test used to determine specific difference – *P*-values - *** < 0.001 and **** < 0.0001.

### *C. difficile* interaction with an established microbiota biofilm

Usually the microbiota, when in a healthy state, would already be established before a *C. difficile* infection. To better replicate this rather than introducing *C. difficile* at the same time as the microbiota, we pre-established the microbiota biofilm for 24 h prior to introducing *C. difficile* (Figure 5A). When comparing this to a *C. difficile* monoculture grown for the same length of time, we observed that having an established microbiota has a significant inhibitory effect on *C. difficile*, both 24 h and 48 h post-addition. The largest difference in *C. difficile* numbers (70-fold) was seen at 24 h post-infection. A pre-established microbiota had a significantly larger impact on *C. difficile* than when the two are seeded together (Figure 5B).

**Figure 5:**
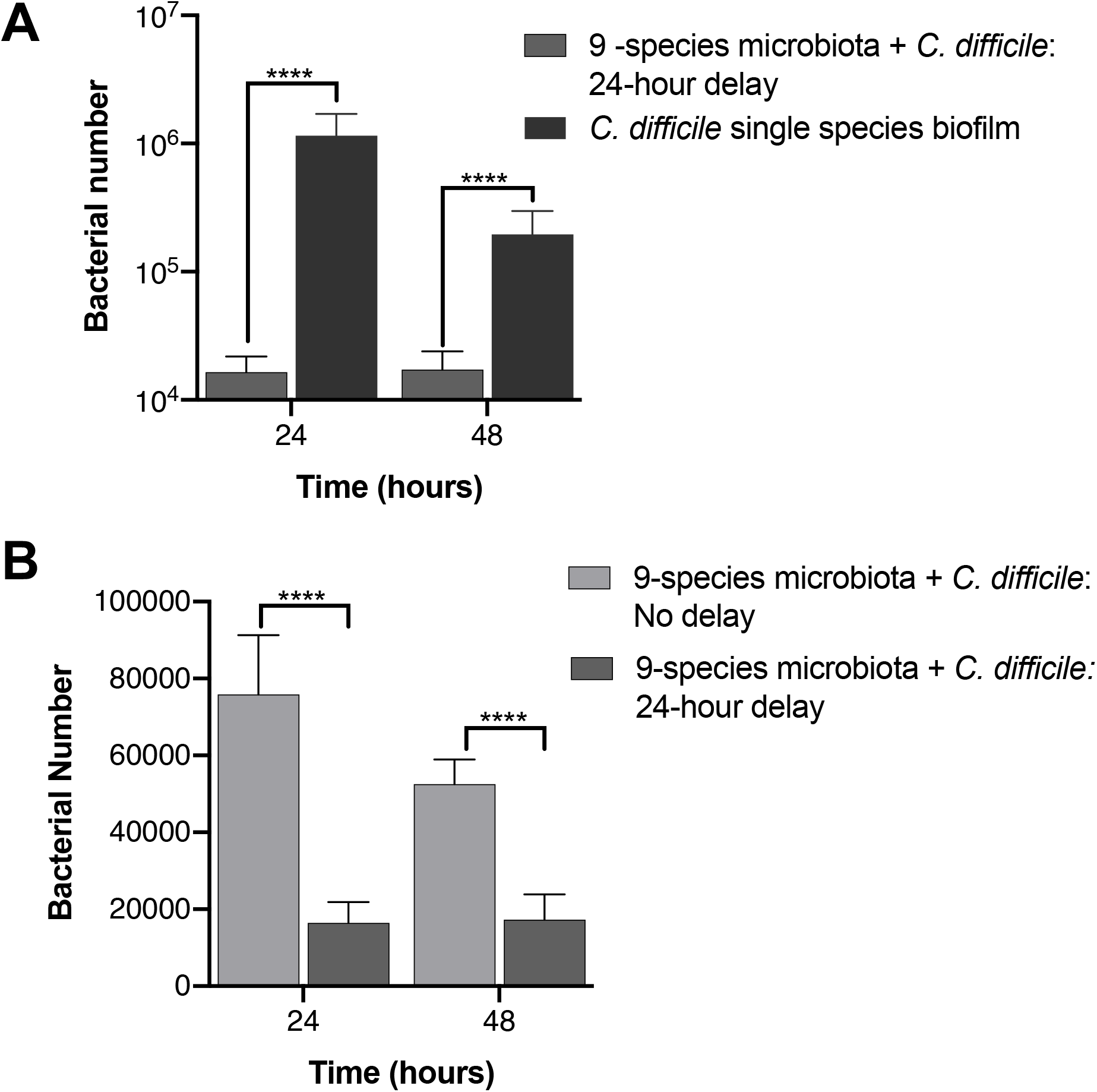
A pre-established microbiota has an inhibitory effect on *C. difficile* growth. **A)** A microbiota biofilm established for 24 h prior to introduction of *C. difficile* (9-species microbiota + *C. difficile* 24 hour delay), was compared to a *C. difficile* only biofilm grown for the same length of time. *C. difficile* numbers were quantified using PMA-qPCR. **B)** The inhibitory effect on *C. difficile* of pre-establishing the microbiota (9-species microbiota + *C. difficile:* 24 hour delay), was compared with the effect on *C. difficile* cocultured with microbiota from the start (9-species microbiota + *C. difficile:* no delay). Data shown are that of three independent experiments in triplicate. An unpaired t-test was used to test for significant difference, *P*-value < 0.0001 - ****.

We tracked each species in the microbiota to examine any changes in response to *C. difficile* (Figures 6 and S5). We expected, as the microbiota was already established, any impact that *C. difficile* had on the microbiota would be lessened compared to when seeded with *C. difficile*. However surprisingly, we found the opposite to be true. Only two species *B. hansenii* and *R. gnavus*, did not change in numbers when compared to a microbiota only control (Supp Figure 6) compared to the four seen when *C. difficile* was added simultaneously (Supp Figure 5). The other seven species, all showed significant differences in numbers when *C. difficile* was introduced (Figure 6). After 24 h with *C. difficile* (48 h from microbiota seeding), we saw a notable positive effect for *B. thetaiotaomicron* and *E. coli*. However, for *F. prausnitzii* we observed a small decrease. At 48 h post-infection with *C. difficile* we still saw a positive effect on *E. coli* but to a much lesser degree. An increase was also observed in *B. ovatus, B. dorei* and *B. adolescentis*. However notably at 48 h, *B. thetaiotaomicron* undergoes a decrease in numbers when compared to the microbiota control, putting it below our detection limit. Thus, although the species impacted by *C. difficile* between simultaneous culture and addition to pre-established microbiota are distinct, there are several species that are affected in both conditions, including the *Bacteroides spp*.

**Figure 1:**
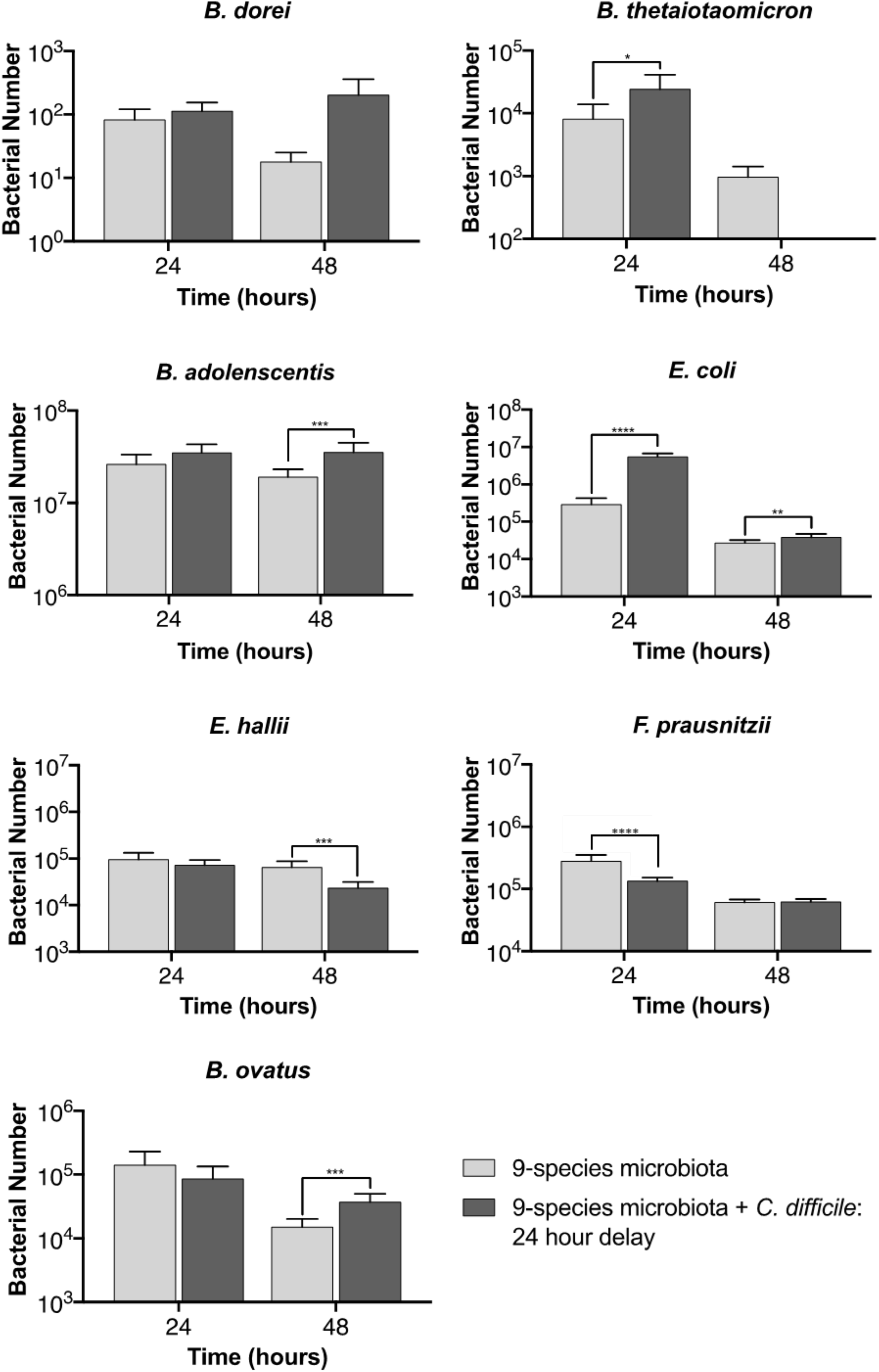
Impact of *C. difficile* on a pre-established microbiota community. PMA-qPCR to track the total number of bacteria for individual species in a nine species representative microbiota biofilm established 24 h before addition of *C. difficile*. Data shown are the mean of three independent experiments in triplicate. Unpaired Student’s t-tests was used to determine significant differences – *P*-values - **** < 0.0001, *** < 0.001, ** < 0.01, * < 0.05.

### Investigating an individual interaction between *Bacteroides* spp and *C. difficile*

We wanted to further investigate if species that were increasing in number alongside the *C. difficile* inhibition, were able to affect *C. difficile* growth. To study if *Bacteroides spp* could impact *C. difficile* growth, we investigated inhibitory effects of *B*. *dorei* when cocultured with *C. difficile* in a dual culture biofilm. Monocultures and a coculture of both species were incubated for 24 h, following determination of CFU counts from the biofilms. In the cocultures, we found that the presence of *B. dorei* significantly reduced (by over 10-fold) the number of *C. difficile* when compared to its monoculture control (Figure 7). In contrast, *B. dorei*, when cultured with *C. difficile* grew far better than when cultured alone. This reduction in *C. difficile* numbers indicated an inhibition mediated by *B. dorei*. To ensure there was no bias in the initial biofilm inoculums, we measured the CFU values of each species, no significant differences were observed between *C. difficile* and *B. dorei* (Supp Figure 7A). Interestingly a similar decrease in bacterial numbers was not observed in planktonic culture of both species (Supp Figure 7B), indicating that the inhibitory effects observed required contact or physical proximity between the two organisms. Our data show that *B. dorei*, one of the commensal *Bacteroides spp* that increases within our complex community in response to addition of *C. difficile*, can negatively impact *C. difficile* growth.

**Figure 2:**
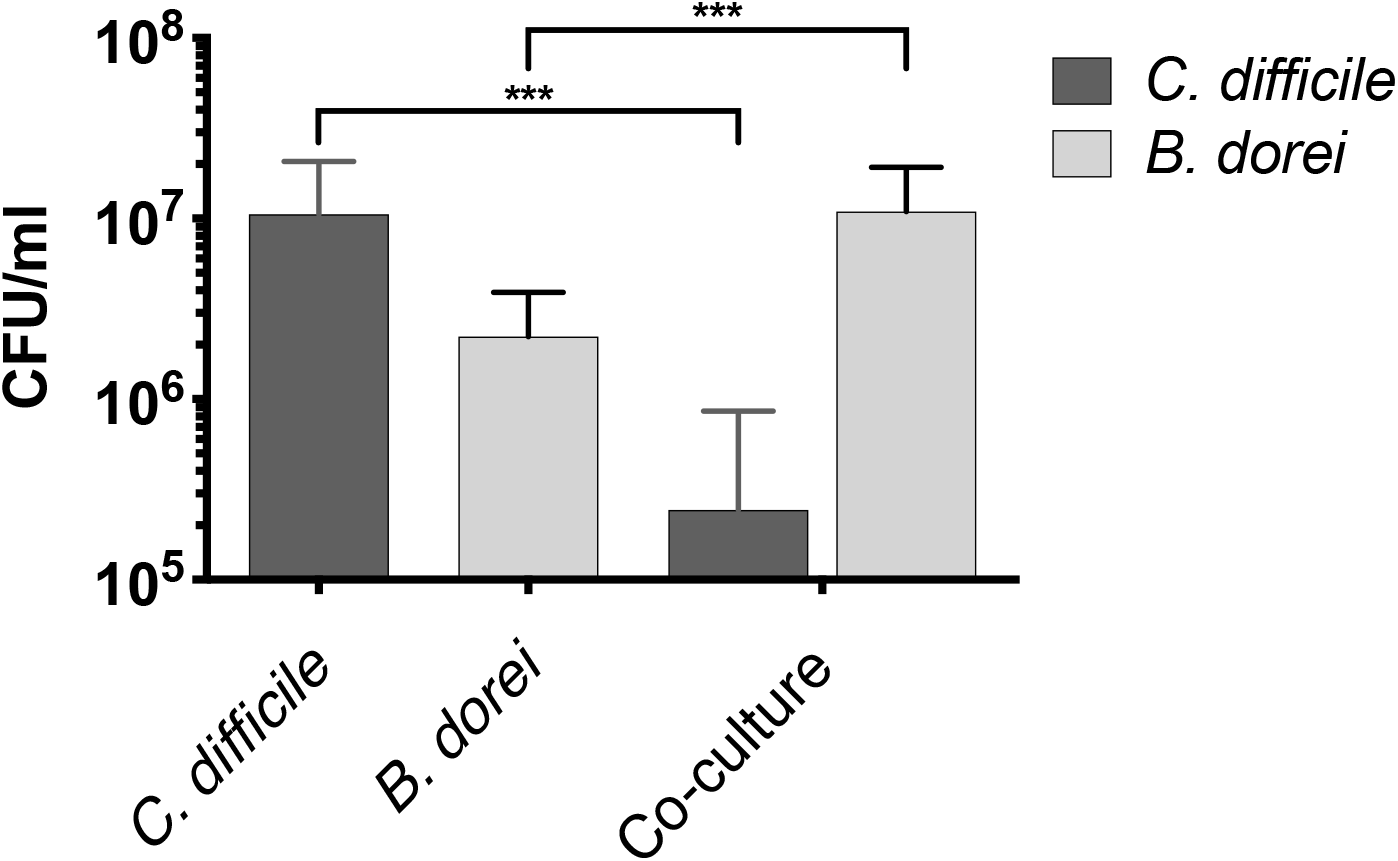
*B. dorei* interactions with *C. difficile* within biofilms. CFU counts of single or mixed cultures of *B. dorei* and *C. difficile* grown on polystyrene plates for 24 h are shown. Data shown are the mean of three independent experiments performed in triplicate, with error bars indicating the standard deviation. Significant difference was determined by an unpaired Students’ t-test, *** indicating a *P*-value < 0.001.

## Discussion

Studying molecular interactions between members of the gut microbiota is important in understanding gut homeostasis and important phenomena like colonisation resistance. We have developed for the first time a complex mixed biofilm community comprising nine representative gut commensal species. We report here a PCR-based method for investigating individual interactions within this complex community. We have quantitated changes in bacterial numbers at different times, and investigated the impact of the gut pathogen *C. difficile* on this community. We demonstrate that several species change in numbers in response to addition of *C. difficile* including *Bacteroides* spp. We have further demonstrated that one of the *Bacteroides* species, *B. dorei* can reduce *C. difficile* growth within dual species biofilms.

Currently, the predominant techniques used to track interaction between species in gut are genomics and metagenomics^3,39–41^. Genome sequencing provides excellent insights into what is present but gives no information on short-term dynamics and underlying bacterial interactions. We chose to use a reductionist approach, coculturing a small number of representative gut species so that the output data was more accessible and increased the resolution at which we could measure changes in individual species. Indeed, we have not covered all the key species but given our chosen quantification technique, it would be possible to now expand this population to include several more species. The main drawback of sequencing is its expense and is hence is usually used for analysis of complex samples which contain hundreds of species. Genome sequencing, like standard qPCR, does not eliminate the quantification of ‘dead’ DNA, which is key in accurate quantification of live bacterial numbers. The other alternative to PMA-qPCR is fluorescent *in situ* hybridization (FISH). Several variations of FISH have been used to quantitate bacteria within communities, however qPCR-based quantitation is generally considered to be more sensitive, rapid and less laborious compared to FISH methods^48–50^. Indeed, FISH as a complementary assay would add key spatial details of the community^31^. Thus, a PMA-qPCR approach is a simple and reliable method to quantitate changes individual members if a complex community.

We were able to successfully track all of the species within a nine species mixed biofilm over a period of 72 h. Interestingly, no single species dominated over others during the experiment, this indicates that rather than direct competition, cross-feeding between species was more likely to be occuring. Henson and Phalak predicted using an *in silico* biofilm metabolic model that a stable cross-feeding relationship between *F. prausnitzii, B. thetaiotaumicron* and *E. coli* could be achieved^51^. However, given the trajectory of several species before the 72 h timepoint we believe this biofilm is as a whole is not stable. The decline in bacterial numbers across all species at 72h, even for species which showed positive signs of growth for the first 48h (such as *B. ovatus* and *B. adolescentis*, Figure 4 & S2) may be due to the spent media becoming toxic and/or growth limiting. Development of a such multibacterial biofilms within flow cells which enable continued flow of nutrients under anaerobic conditions would allow monitoring responses for longer periods of time.

Addition of a pathogen to a commensal community would be expected to induce a response to it. To investigate this, studied responses to the gut pathogen C. difficile was introduced. *C. difficile* is an opportunistic gut pathogen, causing *C. difficile* infection (CDI), which is the leading cause of hospital associated diarrhoea in the US, with half a million new cases each year and a repeat infection rate of 1 in 5 patients^52^. The best defence against *C. difficile* colonisation is a healthy gut microbiota which provides natural immunity to the disease. Antibiotics negatively affect the gut microbiota alongside the pathogens they are aimed at, resulting in alterations of the gut microbiota and an increased likelihood of CDI^19,53^. Distinct changes in the gut microbiota have been associated with CDI^18^, however, as most data are from sequencing faecal samples, information regarding mucosa-associated populations and changes at early points of colonisation. is lacking. Interestingly, the presence of *C. difficile* caused an increase in adhesion for many of the microbiota species within the commensal community (Figure 3A). We predict that *C. difficile* is likely either producing a metabolic bi-product that improves initial growth and binding for species in the microbiota, or that signalling molecules produced by *C. difficile* are detected by the microbiota and inducing a metabolic shift within these species^54^. Further studies using *C. difficile* are necessary to establish whether soluble factors are involved in the increased adherence observed, or whether this is a metabolic effect.

The reduction in the initial adhesion of *C. difficile* observed when cultured with the microbiota (Figure 3A) was expected, as there is a wealth of data on the disruptive effect of the microbiota on *C. difficile* colonisation^8^. The small (approximately 50%) reduction in adherence may be because the microbiota community is not a pre-established, which is normally the case for an infecting *C. difficile*. Bile acids and salts have been shown to have a considerable impact on *C. difficile* ability to colonise the gut, with the microbiota converting the primary bile acids required for *C. difficile* germination such as taurocholic acid^21,55^. The secondary bile acids they convert into can also inhibit *C. difficile* growth^23^. There were no bile acids in the media we use, suggesting that the *C. difficile* inhibition observed was not through the conversion of primary to secondary bile acids. Hence the prevention of bacterial germination through secondary bile acid production appear to be only part of the mechanism for colonisation resistance. Indeed, when the microbiota biofilm was preestablished 24h prior to introducing *C. difficile*, we found that *C. difficile* growth was substantially impacted (Figure 5A & B). Pre-establishing the microbiota could result in increased abundance of the commensal bacteria, and result in higher levels of any secreted inhibitory molecules. Additionally, physical space required for *C. difficile* to adhere would be far less and any nutrients required by *C. difficile* could already be taken out of the media by this stage.

In a successful infection, *C. difficile* has been reported to control the microbiota by modulating bacterial metabolism, including the production of indole^56,57^. It does this through influencing the expression of tryptophanase (tnaA) in other species; *C. difficile* is thought to limit the recovery of the microbiota through indole-mediated inhibition of growth of protective gut bacteria^57^. Small but significant increases in the numbers of *B. dorei, B. ovatus, E. coli, F. prausnitzii*, and *R. gnavus* was observed when *C. difficile* was cocultured with the commensal community. Although of the nine species, *B. ovatus, B. thetaiotaomicron, B. adolescentis, E. coli* and *F. prausnitzii* are indole producers^57–60^, no inhibitory effects were seen. Given that C. *difficile* was instead inhibited, it is possible that the bacteria were unable to reach sufficient numbers to influence indole production. Also, after the addition of *C. difficile* to a preestablished community, we saw an increase in *E. coli* numbers compared to the control (Figure 6). Again, an overabundance of *Proteobacteria* was often found in CDI patients and a characteristic of a successful *C. difficile* infection^20,61^. However, the increase in numbers of multiple *Bacteroides spp* that was observed in parallel may explain the inhibition of *C. difficile* observed in this system. *C. difficile* infections have been associated with a significant decrease in Bacteroidetes, which may suggest a protective role for these bacteria in the gut^61,62^.

Interestingly, *C. difficile* growth is negatively impacted when cocultured with *Bacteroides dorei*, an abundant gut commensal species (Figure 7). A reduction in *C. difficile* growth in presence of *Bacteroides fragilis* was reported previously^63^, appears to be less effective in reducing *C. difficile* growth compared to growth *B. fragilis*. Notably, *B. fragilis* much like *B. dorei* had higher numbers in mixed culture with *C. difficile* than it did on its own and growth inhibitory effects were specific to biofilm growth, indicating that cell-to-cell interactions/ physical proximity may play a role. We also recently reported that *C. difficile* growth on epithelial cells in an *in vitro* gut model was reduced in the presence of *B. doreP^8^. B. fragilis* has been recently reported to prevent *C. difficile* infection in a murine infection model by potentially impacting the integrity of the epithelial barrier^64^. While these data support a role for *Bacteroides spp* in preventing *C. difficile* infection, patients infected with *C. difficile* generally have a reduction in the abundance and diversity of Bacteroidetes^61^. Our data may suggest that without a prior microbiota disturbance, for example with antibiotic treatment, *C. difficile* is unable to bring about a decrease in Bacteroidetes numbers, but instead reinforces Bacteroidetes dominance by improving growth.

In summary, we report a very useful *in vitro* tool that could be used to assess behaviours of members of a complex microbial community. Indeed, this community can be expanded or changed to included other representative species, host cell components (for example, gut epithelial cells) and used to track transcriptomic and metabolic changes modulated in response of stress factors including drugs and pathogens.

## Funding

PhD studentship funded by BBSRC and EPSRC, grant number: EP/L016494/1 to JH. This grant was awarded to the Warwick Integrative Synthetic Biology Centre (Grant Ref: BB/M017982/1) funded under the UK Research Councils’ Synthetic Biology for Growth program.

## Author Contributions

JH performed the experiments for this study. MU and JH were involved in designing experiments in the study. JH and MU wrote and reviewed the manuscript.

## Competing Interests statement

The authors declare no conflict of interest.

## Supplementary Figure legends

**Supplementary Figure 1**. Primer specificity matrix. PCR reactions were conducted with genomic DNA from each individual species to confirm primer specificity. These products were analysed on a 2% agarose gel, with a product size between 100-150 bp depending on the species.

**Supplementary Figure 2**. Standard curves of the nine commensal bacteria and *C. difficile*. Exponentially grown samples of each species were serially diluted to give a range of optical densities. DNA was then extracted from the different dilutions and measured using Qubit and then measured again using qPCR. A semilog regression was fitted to the data. Data shown represent the mean of three technical triplicates.

**Supplementary Figure 3**. A comparison between qPCR predicted bacterial numbers and bacterial numbers obtained from plating (CFU assay). Biofilms of each species were grown for 24 h, these were then washed and resuspended in 1 ml PBS. 50 μl of each sample was taken for serial dilution and plating, the rest was used in the PMA-qPCR pipeline. Data shown represents the mean of three independent experiments in triplicate. A two-way ANOVA showed no significant difference between plating and the qPCR technique.

**Supplementary Figure 4.** Bacterial numbers of individual species in the mixed inoculum used for mixed biofilm assays. Bacterial numbers within the inoculum were tracked using PMA-qPCR. Ct values were converted using our pipeline to represent total bacterial numbers. Data shown is the mean of three independent biological experiments done in in triplicate. Error bars indicate the standard deviation.

**Supplementary Figure 5**: Effects of *C. difficile* on individual species from the 9-species microbiota community. Individual species within a representative microbiota and *C. difficile* biofilm were tracked over 72 h using PMA-qPCR. Data shown are from three independent experiments in triplicate. A two-way ANOVA was used to determine significant differences between the two conditions.

**Supplementary Figure 6**: Effects of *C. difficile* on *R. gnavus* and *B. hansenii* in a pre-established microbiota community. PMA-qPCR was used to track the total numbers of bacteria in a nine species representative microbiota biofilm that was pre-established 24 h before addition of *C. difficile*. Data shown is the mean of three independent experiments in triplicate. Unpaired Student’s t-tests was used to determine presence significant differences.

**Supplementary Figure 7**: **A**) Starting inoculum for *C. difficile – B. dorei* biofilm shows no significant difference when tested with unpaired Student’s t-test. **B**) *B. dorei* and *C. difficile* cultured planktonically as single or mixed cultures for 24 h. Data shown is the mean of three independent tests, with error bars indicating the standard deviation. Significant difference was determined by an unpaired Students t-test, *** indicating a *P*-value < 0.001.

## Supplementary tables

**Table S1**: Primers sequences used in quantification.

**Table S2**: Linearity and primer efficiency of standard curves. Semilog regressions were fitted to each of the standard curves; primer efficiency is calculated from the gradient of these lines.

**Table S3**: Genome length, weight and accession numbers of the sequences used for PMA-qPCR quantification.

## References

1 Guinane, C. M. & Cotter, P. D. Role of the gut microbiota in health and chronic gastrointestinal disease: understanding a hidden metabolic organ. Therap Adv Gastroenterol 6, 295–308, doi:10.1177/1756283X13482996 (2013).

2 Harris, K. G. & Chang, E. B. The intestinal microbiota in the pathogenesis of inflammatory bowel diseases: new insights into complex disease. Clin Sci (Lond) 132, 2013–2028, doi:10.1042/CS20171110 (2018).

3 Qin, J. et al. A metagenome-wide association study of gut microbiota in type 2 diabetes. Nature 490, 55–60, doi:10.1038/nature11450 (2012).

4 Tremaroli, V. & Backhed, F. Functional interactions between the gut microbiota and host metabolism. Nature 489, 242–249, doi:10.1038/nature11552 (2012).

5 Haro, C. et al. Intestinal Microbiota Is Influenced by Gender and Body Mass Index. PLoS One 11, e0154090, doi:10.1371/journal.pone.0154090 (2016).

6 Marchesi, J. R. et al. The gut microbiota and host health: a new clinical frontier. Gut 65, 330–339, doi:10.1136/gutjnl-2015-309990 (2016).

7 Valdes, A. M., Walter, J., Segal, E. & Spector, T. D. Role of the gut microbiota in nutrition and health. BMJ 361, k2179, doi:10.1136/bmj.k2179 (2018).

8 Britton, R. A. & Young, V. B. Role of the intestinal microbiota in resistance to colonization by Clostridium difficile. Gastroenterology 146, 1547–1553, doi:10.1053/j.gastro.2014.01.059 (2014).

9 Lawley, T. D. & Walker, A. W. Intestinal colonization resistance. Immunology 138, 1-11, doi:10.1111/j.1365-2567.2012.03616.x (2013).

10 Sorbara, M. T. & Pamer, E. G. Interbacterial mechanisms of colonization resistance and the strategies pathogens use to overcome them. Mucosal Immunol 12, 1–9, doi:10.1038/s41385-018-0053-0 (2019).

11 Kamada, N. et al. Regulated virulence controls the ability of a pathogen to compete with the gut microbiota. Science 336, 1325–1329, doi:10.1126/science.1222195 (2012).

12 Jacobson, A. et al. A Gut Commensal-Produced Metabolite Mediates Colonization Resistance to Salmonella Infection. Cell Host Microbe 24, 296–307 e297, doi:10.1016/j.chom.2018.07.002 (2018).

13 Willemsen, L. E., Koetsier, M. A., van Deventer, S. J. & van Tol, E. A. Short chain fatty acids stimulate epithelial mucin 2 expression through differential effects on prostaglandin E(1) and E(2) production by intestinal myofibroblasts. Gut 52, 1442–1447, doi:10.1136/gut.52.10.1442 (2003).

14 Zhao, Y. et al. GPR43 mediates microbiota metabolite SCFA regulation of antimicrobial peptide expression in intestinal epithelial cells via activation of mTOR and STAT3. Mucosal Immunol 11, 752–762, doi:10.1038/mi.2017.118 (2018).

15 Saito, K. et al. Inhibition of enterohemorrhagic Escherichia coli O157:H7 infection in a gnotobiotic mouse model with pre-colonization by Bacteroides strains. Biomed Rep 10, 175–182, doi:10.3892/br.2019.1193 (2019).

16 O’Loughlin, J. L. et al. The Intestinal Microbiota Influences Campylobacter jejuni Colonization and Extraintestinal Dissemination in Mice. Appl Environ Microbiol 81, 4642–4650, doi:10.1128/AEM.00281-15 (2015).

17 Leffler, D. A. & Lamont, J. T. Clostridium difficile Infection. N Engl J Med 373, 287–288, doi:10.1056/NEJMc1506004 (2015).

18 Schubert, A. M. et al. Microbiome data distinguish patients with Clostridium difficile infection and non-C. difficile-associated diarrhea from healthy controls. mBio 5, e01021–01014, doi:10.1128/mBio.01021-14 (2014).

19 Schubert, A. M., Sinani, H. & Schloss, P. D. Antibiotic-Induced Alterations of the Murine Gut Microbiota and Subsequent Effects on Colonization Resistance against Clostridium difficile. mBio 6, e00974, doi:10.1128/mBio.00974-15 (2015).

20 Theriot, C. M. & Young, V. B. Interactions Between the Gastrointestinal Microbiome and Clostridium difficile. Annu Rev Microbiol 69, 445–461, doi:10.1146/annurev-micro-091014-104115 (2015).

21 Theriot, C. M., Bowman, A. A. & Young, V. B. Antibiotic-Induced Alterations of the Gut Microbiota Alter Secondary Bile Acid Production and Allow for Clostridium difficile Spore Germination and Outgrowth in the Large Intestine. mSphere 1, doi:10.1128/mSphere.00045-15 (2016).

22 Sorg, J. A. & Sonenshein, A. L. Chenodeoxycholate is an inhibitor of Clostridium difficile spore germination. J Bacteriol 191, 1115–1117, doi:10.1128/JB.01260-08 (2009).

23 Thanissery, R., Winston, J. A. & Theriot, C. M. Inhibition of spore germination, growth, and toxin activity of clinically relevant C. difficile strains by gut microbiota derived secondary bile acids. Anaerobe 45, 86–100, doi:10.1016/j.anaerobe.2017.03.004 (2017).

24 Buffie, C. G. et al. Precision microbiome reconstitution restores bile acid mediated resistance to Clostridium difficile. Nature 517, 205–208, doi:10.1038/nature13828 (2015).

25 Rea, M. C. et al. Thuricin CD, a posttranslationally modified bacteriocin with a narrow spectrum of activity against Clostridium difficile. Proc Natl Acad Sci U S A 107, 9352–9357, doi:10.1073/pnas.0913554107 (2010).

26 Tap, J. et al. Identification of an Intestinal Microbiota Signature Associated With Severity of Irritable Bowel Syndrome. Gastroenterology 152, 111–123 e118, doi:10.1053/j.gastro.2016.09.049 (2017).

27 de Vos, W. M. Microbial biofilms and the human intestinal microbiome. NPJ Biofilms Microbiomes 1, 15005, doi:10.1038/npjbiofilms.2015.5 (2015).

28 Macfarlane, S. & Macfarlane, G. T. Composition and metabolic activities of bacterial biofilms colonizing food residues in the human gut. Appl Environ Microbiol 72, 6204–6211, doi:10.1128/AEM.00754-06 (2006).

29 Vandeplassche, E., Coenye, T. & Crabbe, A. Developing selective media for quantification of multispecies biofilms following antibiotic treatment. PLoS One 12, e0187540, doi:10.1371/journal.pone.0187540 (2017).

30 Klein, M. I., Scott-Anne, K. M., Gregoire, S., Rosalen, P. L. & Koo, H. Molecular approaches for viable bacterial population and transcriptional analyses in a rodent model of dental caries. Mol Oral Microbiol 27, 350–361, doi:10.1111/j.2041-1014.2012.00647.x (2012).

31 Ammann, T. W., Bostanci, N., Belibasakis, G. N. & Thurnheer, T. Validation of a quantitative real-time PCR assay and comparison with fluorescence microscopy and selective agar plate counting for species-specific quantification of an in vitro subgingival biofilm model. J Periodontal Res 48, 517–526, doi:10.1111/jre.12034 (2013).

32 Loozen, G., Boon, N., Pauwels, M., Quirynen, M. & Teughels, W. Live/dead real-time polymerase chain reaction to assess new therapies against dental plaque-related pathologies. Mol Oral Microbiol 26, 253–261, doi:10.1111/j.2041-1014.2011.00615.x (2011).

33 Yasunaga, A. et al. Monitoring the prevalence of viable and dead cariogenic bacteria in oral specimens and in vitro biofilms by qPCR combined with propidium monoazide. BMC Microbiol 13, 157, doi:10.1186/1471-2180-13-157 (2013).

34 Haarman, M. & Knol, J. Quantitative real-time PCR assays to identify and quantify fecal Bifidobacterium species in infants receiving a prebiotic infant formula. Appl Environ Microbiol 71, 2318–2324, doi:10.1128/AEM.71.5.2318-2324.2005 (2005).

35 Matsuki, T., Watanabe, K., Fujimoto, J., Takada, T. & Tanaka, R. Use of 16S rRNA gene-targeted group-specific primers for real-time PCR analysis of predominant bacteria in human feces. Appl Environ Microbiol 70, 7220–7228, doi:10.1128/AEM.70.12.7220-7228.2004 (2004).

36 Ye, J. et al. Primer-BLAST: a tool to design target-specific primers for polymerase chain reaction. BMC Bioinformatics 13, 134, doi:10.1186/1471-2105-13-134 (2012).

37 Huang, R., Zhang, J., Yang, X. F. & Gregory, R. L. PCR-Based Multiple Species Cell Counting for In Vitro Mixed Culture. PLoS One 10, e0126628, doi:10.1371/journal.pone.0126628 (2015).

38 Anonye, B. O. et al. Probing Clostridium difficile Infection in Complex Human Gut Cellular Models. Front Microbiol 10, 879, doi:10.3389/fmicb.2019.00879 (2019).

39 Qin, J. et al. A human gut microbial gene catalogue established by metagenomic sequencing. Nature 464, 59–65, doi:10.1038/nature08821 (2010).

40 Li, J. et al. An integrated catalog of reference genes in the human gut microbiome. Nat Biotechnol 32, 834–841, doi:10.1038/nbt.2942 (2014).

41 Schloissnig, S. et al. Genomic variation landscape of the human gut microbiome. Nature 493, 45–50, doi:10.1038/nature11711 (2013).

42 Eckburg, P. B. et al. Diversity of the human intestinal microbial flora. Science 308, 1635–1638, doi:10.1126/science.1110591 (2005).

43 Nocker, A., Sossa-Fernandez, P., Burr, M. D. & Camper, A. K. Use of propidium monoazide for live/dead distinction in microbial ecology. Appl Environ Microbiol 73, 5111–5117, doi:10.1128/AEM.02987-06 (2007).

44 Yang, X., Badoni, M. & Gill, C. O. Use of propidium monoazide and quantitative PCR for differentiation of viable Escherichia coli from E. coli killed by mild or pasteurizing heat treatments. Food Microbiol 28, 1478–1482, doi:10.1016/j.fm.2011.08.013 (2011).

45 Lee, E. S., Lee, M. H. & Kim, B. S. Evaluation of propidium monoazide-quantitative PCR to detect viable Mycobacterium fortuitum after chlorine, ozone, and ultraviolet disinfection. Int J Food Microbiol 210, 143–148, doi:10.1016/j.ijfoodmicro.2015.06.019 (2015).

46 Tavernier, S. & Coenye, T. Quantification of Pseudomonas aeruginosa in multispecies biofilms using PMA-qPCR. PeerJ 3, e787, doi:10.7717/peerj.787 (2015).

47 Svec, D., Tichopad, A., Novosadova, V., Pfaffl, M. W. & Kubista, M. How good is a PCR efficiency estimate: Recommendations for precise and robust qPCR efficiency assessments. Biomol Detect Quantif 3, 9–16, doi:10.1016/j.bdq.2015.01.005 (2015).

48 Amann, R. I., Krumholz, L. & Stahl, D. A. Fluorescent-oligonucleotide probing of whole cells for determinative, phylogenetic, and environmental studies in microbiology. J Bacteriol 172, 762–770, doi:10.1128/jb.172.2.762-770.1990 (1990).

49 Rigottier-Gois, L., Bourhis, A. G., Gramet, G., Rochet, V. & Dore, J. Fluorescent hybridisation combined with flow cytometry and hybridisation of total RNA to analyse the composition of microbial communities in human faeces using 16S rRNA probes. FEMS Microbiol Ecol 43, 237–245, doi:10.1111/j.1574-6941.2003.tb01063.x (2003).

50 Cleusix, V., Lacroix, C., Dasen, G., Leo, M. & Le Blay, G. Comparative study of a new quantitative real-time PCR targeting the xylulose-5-phosphate/fructose-6-phosphate phosphoketolase bifidobacterial gene (xfp) in faecal samples with two fluorescence in situ hybridization methods. J Appl Microbiol 108, 181–193, doi:10.1111/j.1365-2672.2009.04408.x (2010).

51 Henson, M. A. & Phalak, P. Microbiota dysbiosis in inflammatory bowel diseases: in silico investigation of the oxygen hypothesis. BMC Syst Biol 11, 145, doi:10.1186/s12918-017-0522-1 (2017).

52 Lessa, F. C., Winston, L. G., McDonald, L. C. & Emerging Infections Program, C. d. S. T. Burden of Clostridium difficile infection in the United States. N Engl J Med 372, 2369–2370, doi:10.1056/NEJMc1505190 (2015).

53 Haak, B. W. et al. Long-term impact of oral vancomycin, ciprofloxacin and metronidazole on the gut microbiota in healthy humans. J Antimicrob Chemother 74, 782–786, doi:10.1093/jac/dky471 (2019).

54 Fuentes, S. et al. Reset of a critically disturbed microbial ecosystem: faecal transplant in recurrent Clostridium difficile infection. ISME J 8, 1621–1633, doi:10.1038/ismej.2014.13 (2014).

55 Winston, J. A. & Theriot, C. M. Impact of microbial derived secondary bile acids on colonization resistance against Clostridium difficile in the gastrointestinal tract. Anaerobe 41, 44–50, doi:10.1016/j.anaerobe.2016.05.003 (2016).

56 Jenior, M. L., Leslie, J. L., Young, V. B. & Schloss, P. D. Clostridium difficile Alters the Structure and Metabolism of Distinct Cecal Microbiomes during Initial Infection To Promote Sustained Colonization. mSphere 3, doi:10.1128/mSphere.00261-18 (2018).

57 Darkoh, C., Plants-Paris, K., Bishoff, D. & DuPont, H. L. Clostridium difficile Modulates the Gut Microbiota by Inducing the Production of Indole, an Interkingdom Signaling and Antimicrobial Molecule. mSystems 4, doi:10.1128/mSystems.00346-18 (2019).

58 Whaley, D. N., Wiggs, L. S., Miller, P. H., Srivastava, P. U. & Miller, J. M. Use of Presumpto Plates to identify anaerobic bacteria. J Clin Microbiol 33, 1196–1202 (1995).

59 Aragozzini, F., Ferrari, A., Pacini, N. & Gualandris, R. Indole-3-lactic acid as a tryptophan metabolite produced by Bifidobacterium spp. Appl Environ Microbiol 38, 544–546 (1979).

60 Berstad, A., Raa, J. & Valeur, J. Indole - the scent of a healthy ‘inner soil’. Microb Ecol Health Dis 26, 27997, doi:10.3402/mehd.v26.27997 (2015).

61 Reeves, A. E. et al. The interplay between microbiome dynamics and pathogen dynamics in a murine model of Clostridium difficile Infection. Gut Microbes 2, 145–158, doi:10.4161/gmic.2.3.16333 (2011).

62 Peterfreund, G. L. et al. Succession in the gut microbiome following antibiotic and antibody therapies for Clostridium difficile. PLoS One 7, e46966, doi:10.1371/journal.pone.0046966 (2012).

63 Slater, R. T., Frost, L. R., Jossi, S. E., Millard, A. D. & Unnikrishnan, M. Clostridioides difficile LuxS mediates inter-bacterial interactions within biofilms. Sci Rep 9, 9903, doi:10.1038/s41598-019-46143-6 (2019).

64 Deng, H. et al. Bacteroides fragilis Prevents Clostridium difficile Infection in a Mouse Model by Restoring Gut Barrier and Microbiome Regulation. Front Microbiol 9, 2976, doi:10.3389/fmicb.2018.02976 (2018).

