## supplemental figures for "Dissecting individual pathogen-commensal interactions within a complex gut microbiota community"

### Supp Figure 1

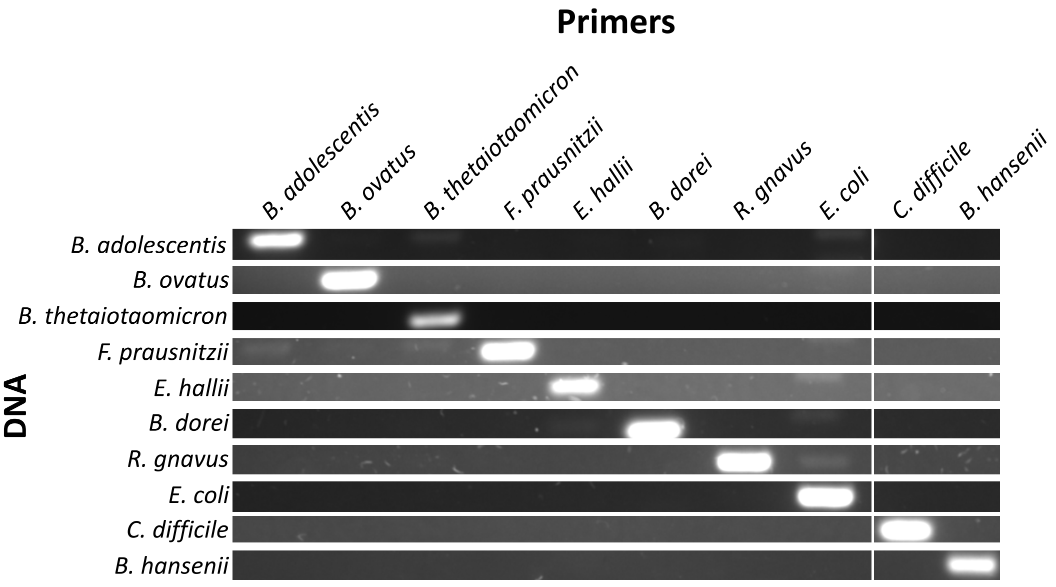

### Supp Figure 2

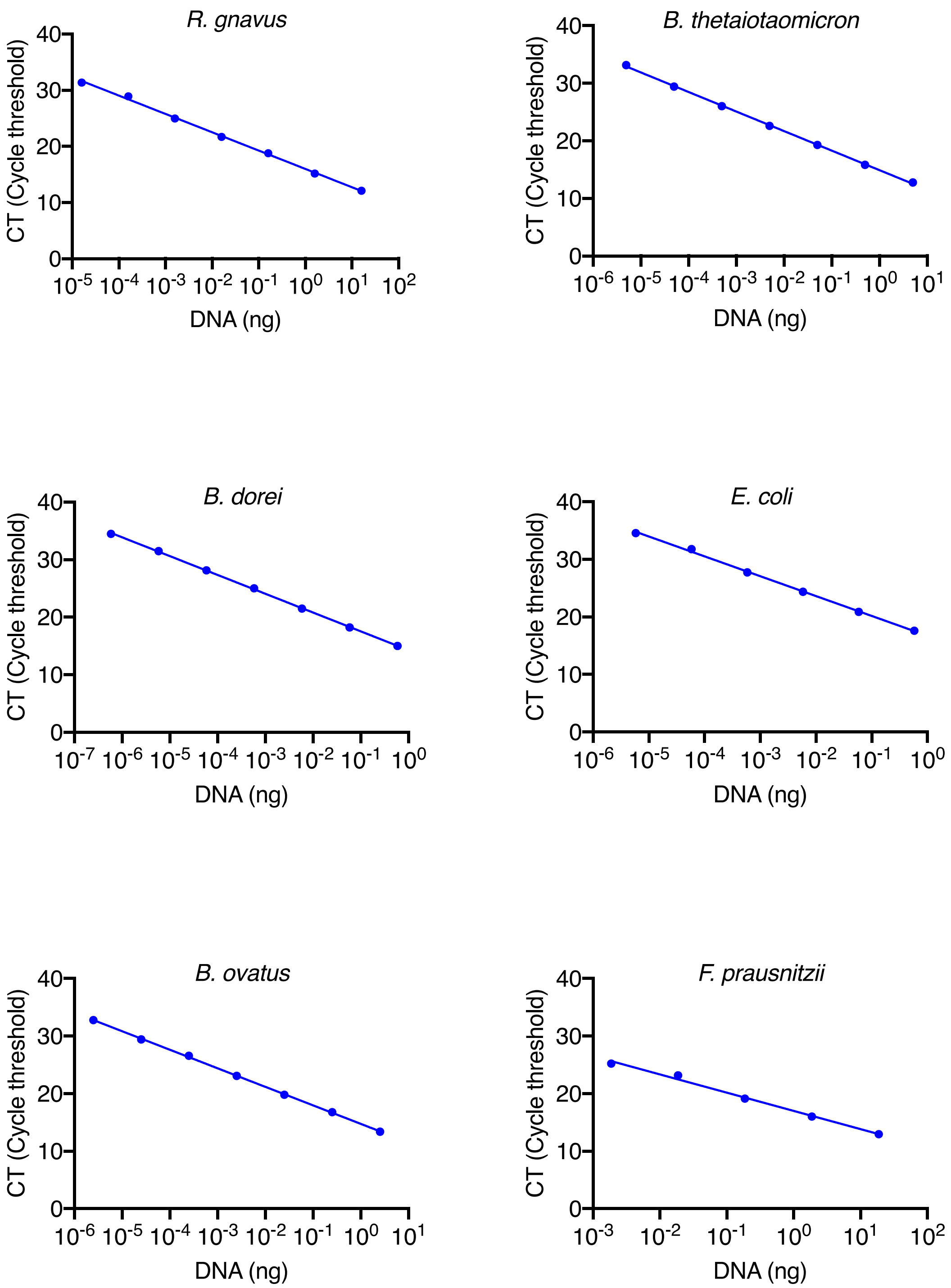

### Supp Figure 2 (continued)

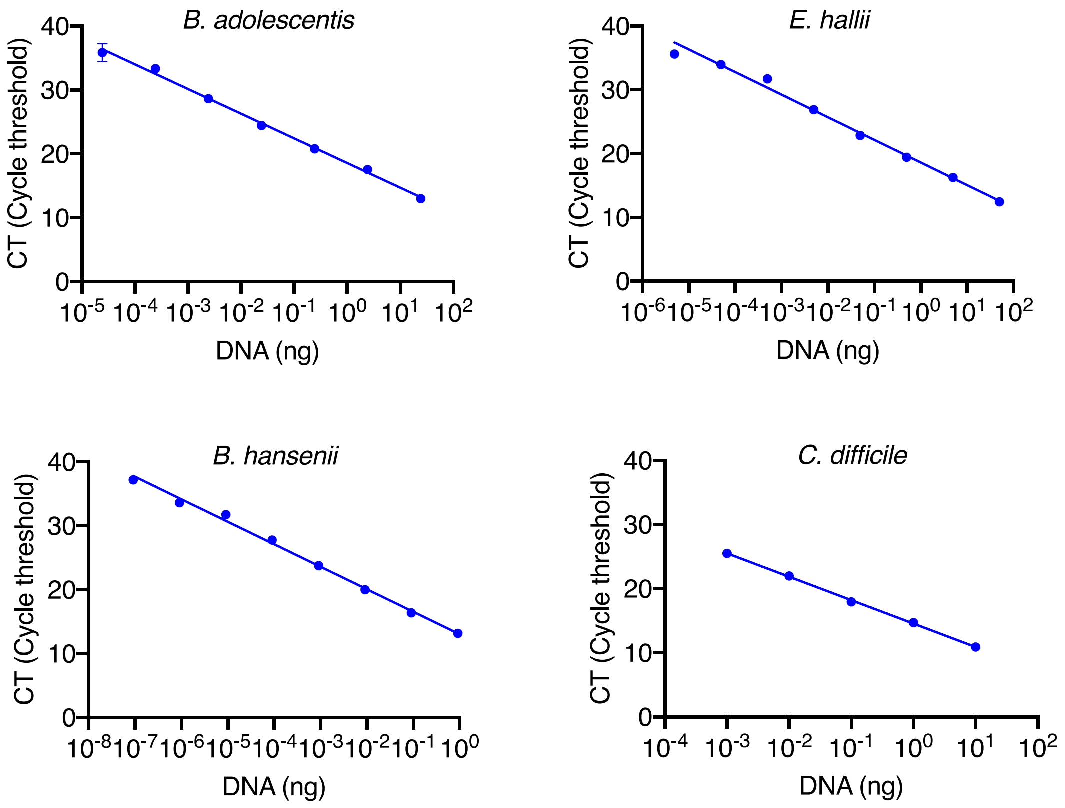

Supp Figure 3

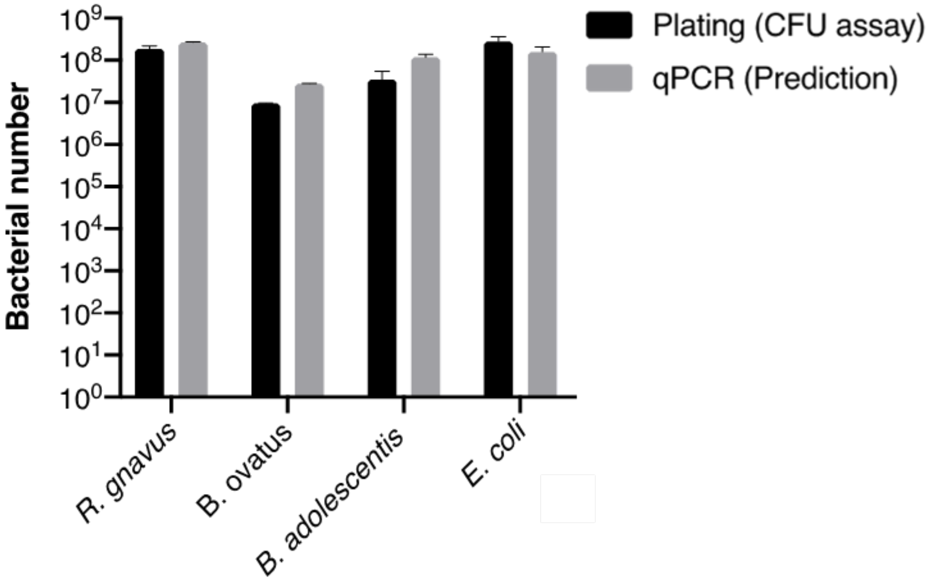

Supp Figure 4

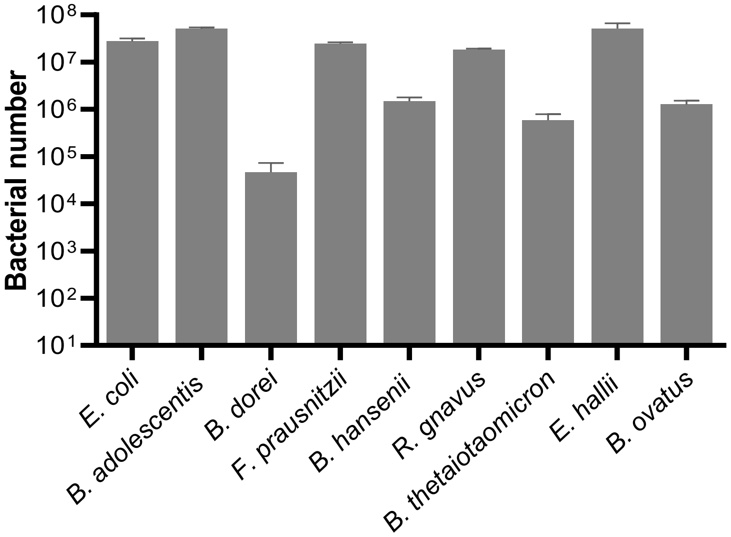

### Supp Figure 5

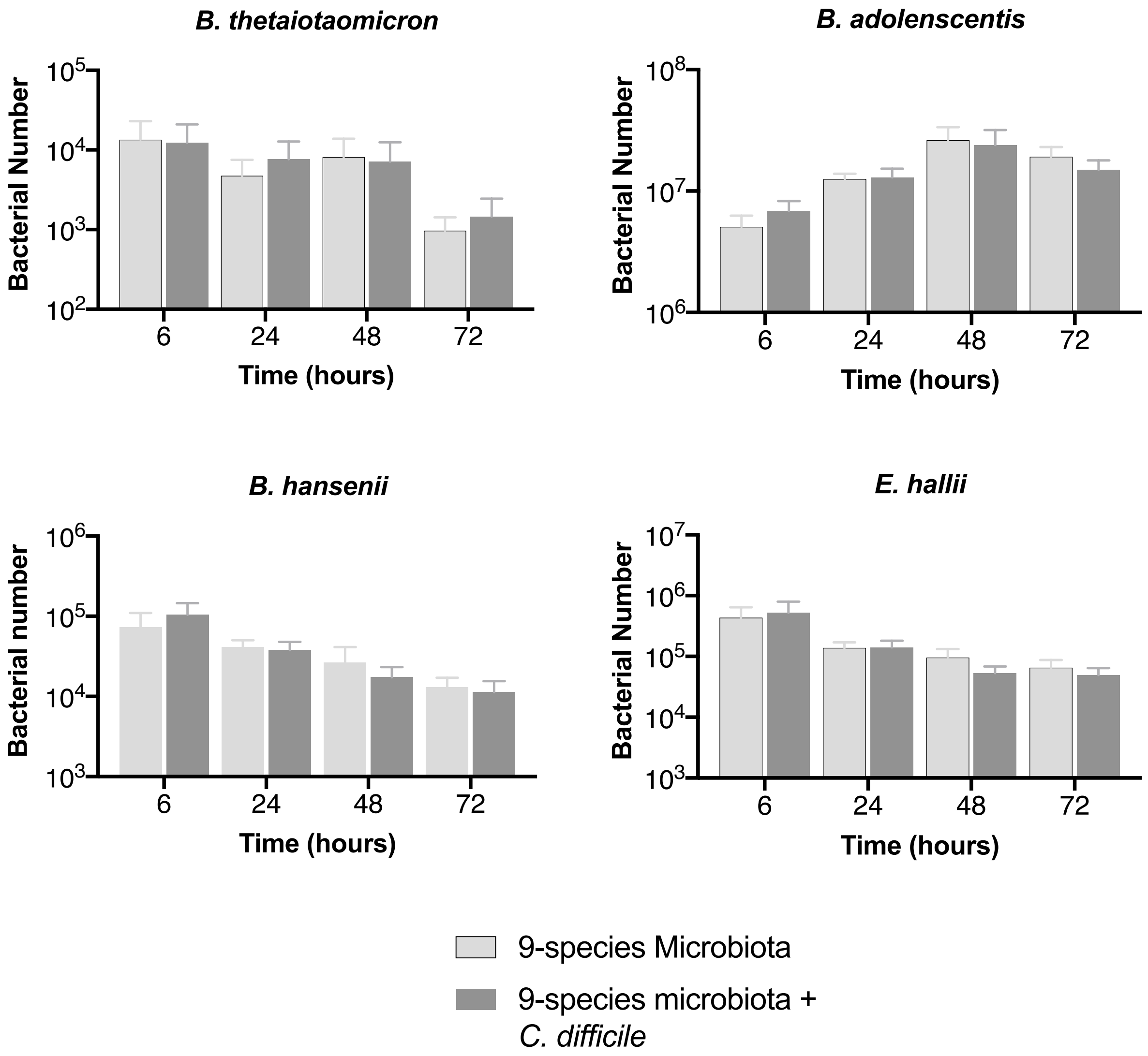

Supp Figure 6

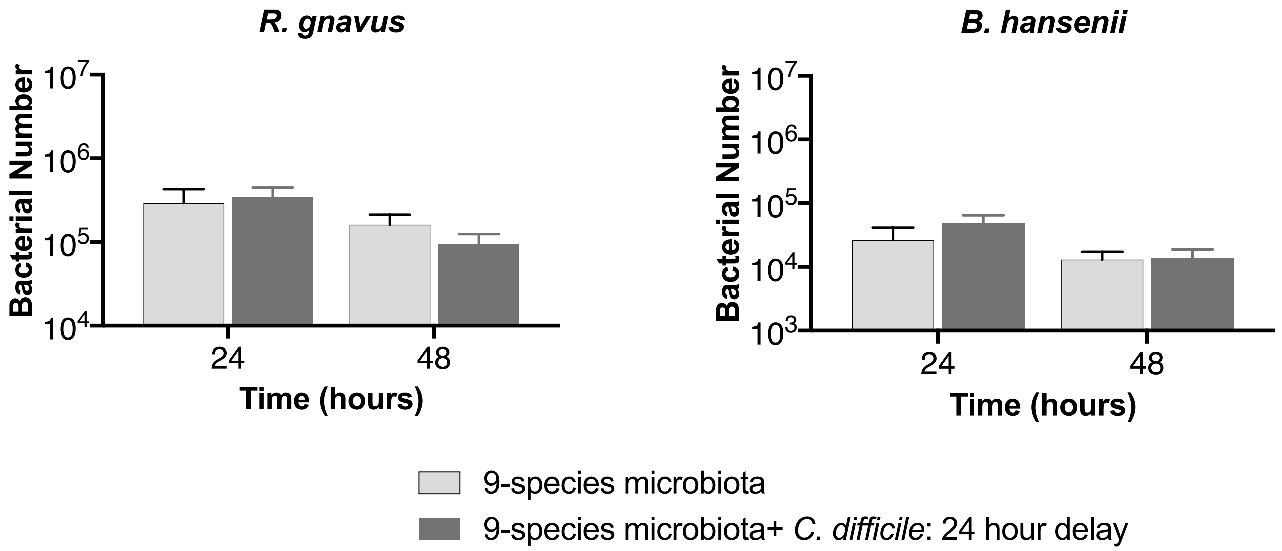

Supp Figure 7

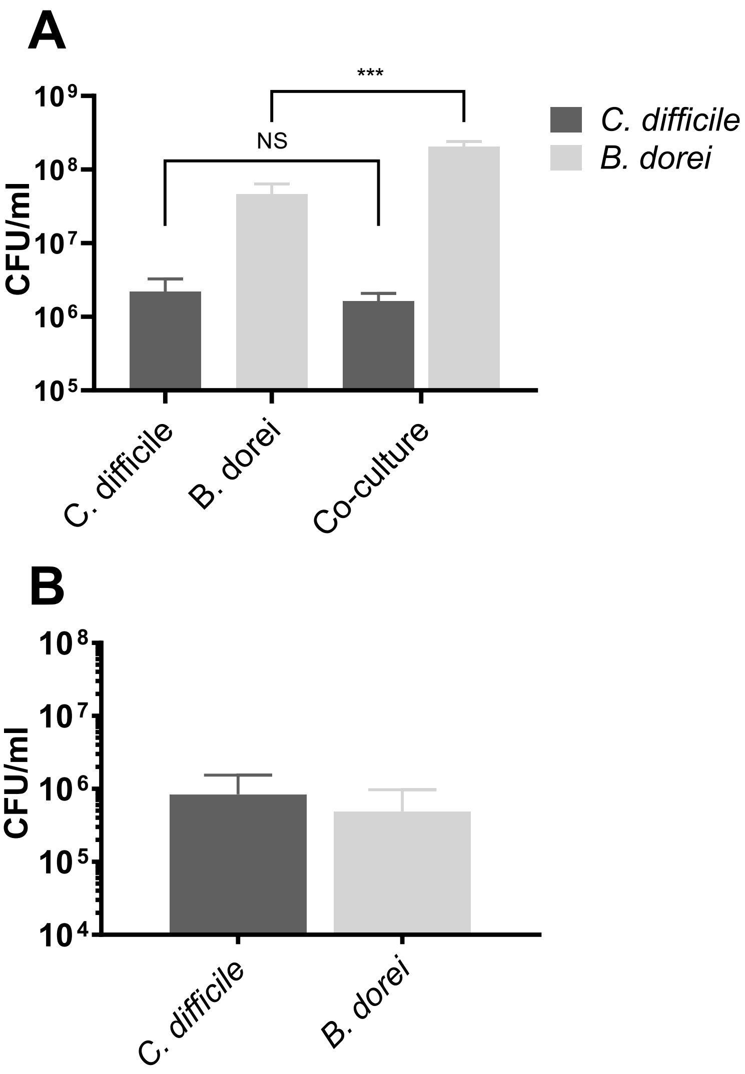
