## Supplementary material for "Dissecting individual pathogen-commensal interactions within a complex gut microbiota community": supplmental tables

### Table S1

| Species | Gene of interest | Sequences |
|---|---|---|
| *B. dorei* | *topI* | Forward: AAGCGGCTTCAAGAAACAGG<br>Reverse: GTGCCCTTTACCTTGGGAAC |
| *B. ovatus* | *topI* | Forward: GGGCCTATTATCGCAACCGA<br>Reverse: AGGTGCATACGTAGACGGAC |
| *B. thetaiotaomicron* | *topI* | Forward: GTCTGTAATCAAGTCCGCCG<br>Reverse: AATGCCGGAAAGCGGTAAAC |
| *B. adolescentis* | *topI* | Forward: CTCCGGATACACGGTCATGG<br>Reverse: GTCTTCGATATCCACGCCGA |
| *B. hansenii* | *gyrA* | Forward: GACGTAAGAAGCACCGGTAGA<br>Reverse: ATAATCGCCCTGACAGGTAAGC |
| *C. difficile* | *gyrA* | Forward: GGTTGAAAGAATAGCAGAGTTAGTT<br>Reverse: GCATTAGCATCCCTCTTTAATTCTA |
| *E. coli* | *gyrA* | Forward: GAACTCGGTGAGGACGGTTT<br>Reverse: GCTGGAACAGGACGAACGTA |
| *E. hallii* | *gyrA* | Forward: TACCGCCTCATCGGACTTGA<br>Reverse: TCATGGAGGCTGGATGCTCT |
| *F. prausnitzii* | *gyrA* | Forward: CCGGTGTCCGTGTCATGC<br>Reverse: CTCAGCCTCTACTGTCTCGG |
| *R. gnavus* | *gyrA* | Forward: GCTGAACAGAGCAGAAGAGC<br>Reverse: TCCTTCGCAGTCTGAACATTCT |

### Table S2

| Species | Slope | Y-intercept | $R^2$ | Primer efficiency (%) |
|---|---|---|---|---|
| *B. dorei* | -3.269 | 14.26 | 0.9994 | 102.26 |
| *B. ovatus* | -3.224 | 14.74 | 0.9994 | 104.26 |
| *B. thetaiotaomicron* | -3.396 | 14.91 | 0.9992 | 97.00 |
| *B. adolescentis* | -3.858 | 18.57 | 0.9944 | 81.64 |
| *B. hansenii* | -3.522 | 13.03 | 0.9951 | 92.28 |
| *C. difficile* | -3.651 | 14.56 | 0.9991 | 87.89 |
| *E. coli* | -3.450 | 16.73 | 0.9986 | 94.92 |
| *E. hallii* | -3.545 | 18.61 | 0.9903 | 91.46 |
| *F. prausnitzii* | -3.171 | 17.01 | 0.9927 | 106.71 |
| *R. gnavus* | -3.263 | 15.98 | 0.9979 | 102.52 |

### Table S3

| Species | Length (b) | Weight (ng) | NCBI Reference |
|---|---|---|---|
| *B. dorei* | 6079576 | $6.56 \times 10^{-6}$ | GCF_000273035.1 |
| *B. ovatus* | 6549476 | $7.07 \times 10^{-6}$ | GCF_000218325.1 |
| *B. thetaiotaomicron* | 6293399 | $6.79 \times 10^{-6}$ | GCF_000011065.1 |
| *B. adolescentis* | 2389110 | $2.58 \times 10^{-6}$ | GCF_000154085.1 |
| *B. hansenii* | 3065949 | $3.31 \times 10^{-6}$ | GCF_002222595.2 |
| *C. difficile* | 4191339 | $4.52 \times 10^{-6}$ | GCF_000027105.1 |
| *E. coli* | 5106156 | $5.51 \times 10^{-6}$ | GCF_000159295.1 |
| *E. hallii* | 3290996 | $3.55 \times 10^{-6}$ | GCF_000173975.1 |
| *F. prausnitzii* | 3090349 | $3.34 \times 10^{-6}$ | GCF_000162015.1 |
| *R. gnavus* | 3181861 | $3.43 \times 10^{-6}$ | GCF_000507805.1 |
